# Meta-analysis of metagenomes via machine learning and assembly graphs reveals strain switches in Crohn’s disease

**DOI:** 10.1101/2022.06.30.498290

**Authors:** Taylor E. Reiter, Luiz Irber, Alicia A. Gingrich, Dylan Haynes, N. Tessa Pierce-Ward, Phillip T. Brooks, Yosuke Mizutani, Dominik Moritz, Felix Reidl, Amy D. Willis, Blair D. Sullivan, C. Titus Brown

## Abstract

Microbial strains have closely related genomes but may have different phenotypes in the same environment. Shotgun metagenomic sequencing can capture the genomes of all strains present in a community but strain-resolved analysis from shotgun sequencing alone remains difficult. We developed an approach to identify and interrogate strain-level differences in groups of metagenomes. We use this approach to perform a meta-analysis of stool microbiomes from individuals with and without inflammatory bowel disease (IBD; Crohn’s disease, ulcerative colitis; n = 605), a disease for which there are not specific microbial biomarkers but some evidence that microbial strain variation may stratify by disease state. We first developed a machine learning classifier based on compressed representations of complete metagenomes (FracMinHash sketches) and identified genomes that correlate with IBD subtype. To rescue variation that may not have been present in the genomes, we then used assembly graph genome queries to recover strain variation for correlated genomes. Lastly, we developed a novel differential abundance framework that works directly on the assembly graph to uncover all sequence variants correlated with IBD. We refer to this approach as dominating set differential abundance analysis and have implemented it in the [spacegraphcats software package](https://github.com/spacegraphcats/spacegraphcats). Using this approach, we identified five bacterial strains that are associated with Crohn’s disease. Our method captures variation within the entire sequencing data set, allowing for discovery of previously hidden disease associations.

## Introduction

Sub-species groupings of microorganisms have functional differences that govern important genome-environment interactions across diverse ecosystems. For example, ecotypes of *Escherichia coli* have different gene complements that allow each group to thrive in diverse environments like the gut, soil, and freshwater [1]. Metagenomic sequencing data from communities of microorganisms contain information about specific strains present in a sample, but strain-resolved insights are lacking due to incomplete references or inability of current tools to retrieve such information [2]. Here we use *strain* to refer to within-species variation that generates taxonomic grouping below the species level.

Inflammatory bowel disease (IBD) is a group of disorders that are characterized by chronic inflammation of the intestines which may in part be the result of host-mediated inflammatory responses to microorganisms [3]. IBD classically manifests in three subtypes depending on clinical presentation, including Crohn’s disease (CD), which presents as discontinuous patches of inflammation throughout the gastrointestinal tract, ulcerative colitis (UC), which presents as continuous inflammation isolated to the colon and rectum, and undetermined subtype, which cannot be clinically or biologically distinguished as CD or UC. Diagnosis can be clinically difficult, with ramifications associated with incorrect treatment resulting in unnecessary patient morbidity. Detection of microbial signatures associated with IBD subtype may lead to improved diagnostic criteria and therapeutics that extend periods of remission. However, such signatures have thus far remained elusive [4].

The microbiome of CD and UC is heterogeneous, and studies that characterize the microbiome often produce conflicting results [4]. This is likely in part driven by large inter- and intra-individual variation [5], but is also attributable to non-standardized laboratory, sequencing, and analysis techniques used to profile the gut microbiome [4]. Dysbiosis is frequently observed in IBD, particularly in CD [6,7,8,9,10], however dysbiosis alone is not a signature of IBD [5]. *Dysbiosis* is defined as a decrease in gut microbial diversity that results in an imbalance between protective and harmful microorganisms, leading to intestinal inflammation [11].

Strain-level differences may account for some heterogeneity in IBD gut microbiome profiles. A recent investigation of time-series gut microbiome metagenomes found that one clade of *Ruminococcus gnavus* is enriched in CD [12]. Further, this clade produces an inflammatory polysaccharide [13,14]. While this clade is enriched in CD, its enrichment was previously masked from computational discovery by concomitant decreases in other *Ruminococcus* species in IBD [12], highlighting the need for strain-resolved analysis of metagenomic sequencing in the exploration of IBD gut microbiomes.

Given these features of the IBD gut microbiome, strain-resolved analysis may yield insights into the dynamics of these communities. The two biggest obstacles to strain-level analysis of short read data are the lack of strain representation in databases together with the challenge of haplotype-level resolution in assembly and binning. While long reads have made strides toward resolving the latter issue [15], in habitats like the gut where communities are dominated by single strains of microbes [16] the largest barrier to strain-level analysis is the exclusion of data that does not match to reference databases. New data analysis techniques are needed to make full use of strain level data.

*K-mers*, words of length *k* in nucleotide sequences, have previously been used for annotation-free characterization of sequencing data [17,18,19]. K-mers are suitable for strain-resolved metagenome analysis because their absence in reference databases does not preclude their analysis. Moreover, k-mer analysis does not rely on marker genes which are largely conserved at the strain level, and k-mers are suitable for species- and strain-level classification [20,21]. Investigating all k-mers in metagenomes is more computationally intensive than reference-based approaches [22], but data-reduction techniques like FracMinHash sketching make k-mer-based analysis scalable to large-scale sequence comparisons [23,24]. FracMinHash sketching sacrifices the fine-scaled resolution of reference-based techniques but is representative of the full sequencing sample and can make use of all available genomes [21], thus including strain-variable accessory elements that may be associated with diseases

Like sketches, assembly graphs also represent k-mers in a metagenome, but assembly graphs retain important sequencing context and can aggregate known functional and taxonomic annotations, recovering critical information lost through sketching approaches [25,26]. While assembly graphs have been leveraged in metagenome analyses [28], their large size precludes analysis at scale. The *spacegraphcats* tool is designed to tackle this issue, implementing algorithms that efficiently reduce the size of an assembly graph and enabling rapid querying and sequence retrieval [25]. These algorithms center around dominating sets, which partition the graph into *pieces* by assigning every node to a graph-localized neighborhood [25]. This simplified graph enables efficient queries: querying with a sequence that overlaps any k-mer in a compact de Bruijn graph (cDBG) node returns all k-mers (or all reads containing those k-mers) from the graph neighborhood. Genome queries often recover sequences not in reference databases or *de novo* assemblies, which disproportionately include sequences from both low coverage regions and highly variable portions of the graph (e.g. sequencing reads that neither assemble nor bin) [25]. When a query has a containment index between 10^-2^ and 10^-3^ with the assembly graph, 20-40% of a target genome sequence is recovered from a metagenome query, and for containment indices above 10^-1^ this increases to >80% [25]. Containment index is calculated by comparing the relative size of the intersection to the union between k-mers in a query and k-mers in a metagenome [29].

Here, we develop k-mer- and assembly graph-based techniques to perform a meta-analysis of stool microbiome metagenomes from individuals with (CD, UC) and without (nonIBD) IBD [5,8,10,12,30,31]. Using these approaches, we detect a consistent signature of IBD subtype in fecal microbiome metagenomes. We identify a small set of k-mers that are predictive of UC and CD, and find that these k-mers originate from a core set of microbial genomes. We find that a stochastic loss of diversity in this core set of microbial genomes is a hallmark of CD, and to a lesser extent, UC, as has been previously demonstrated [4]. While reduced diversity is responsible for the majority of disease signatures, we also find signatures of strains present in the disease state. Sequences associated with these strains occurred more frequently in IBD metagenomes but are present in low abundance in nonIBD metagenomes as well. Our approach provides a solution for strain-level analysis of short read metagenomic data sets, and our findings provide future avenues for research into IBD therapeutics.

## Results

We developed a computational approach to resolve sub-species level differences between groups of short read shotgun metagenomes (**Figure 1**). While our pipeline relies on many published algorithms, we developed two new approaches that, when combined with existing tools, generated insights into microbial sequences associated with IBD subtype. After consistent pre-processing, we used FracMinHash sketching to produce subsampled k-mer abundance profiles of metagenomes that reflected the sequence diversity in a sample [21,23], and used these profiles to perform metagenome-wide k-mer association with IBD subtype. We refer to FracMinHash sketches as *sketches* or k-mer abundance profiles, and for simplicity, continue referring to the sub-sampled k-mers in a sketch as *k-mers*. Retaining only k-mers associated with IBD, we used a minimum set cover approach to identify the genomes that best encompassed these k-mers [21].

**Figure 1:**
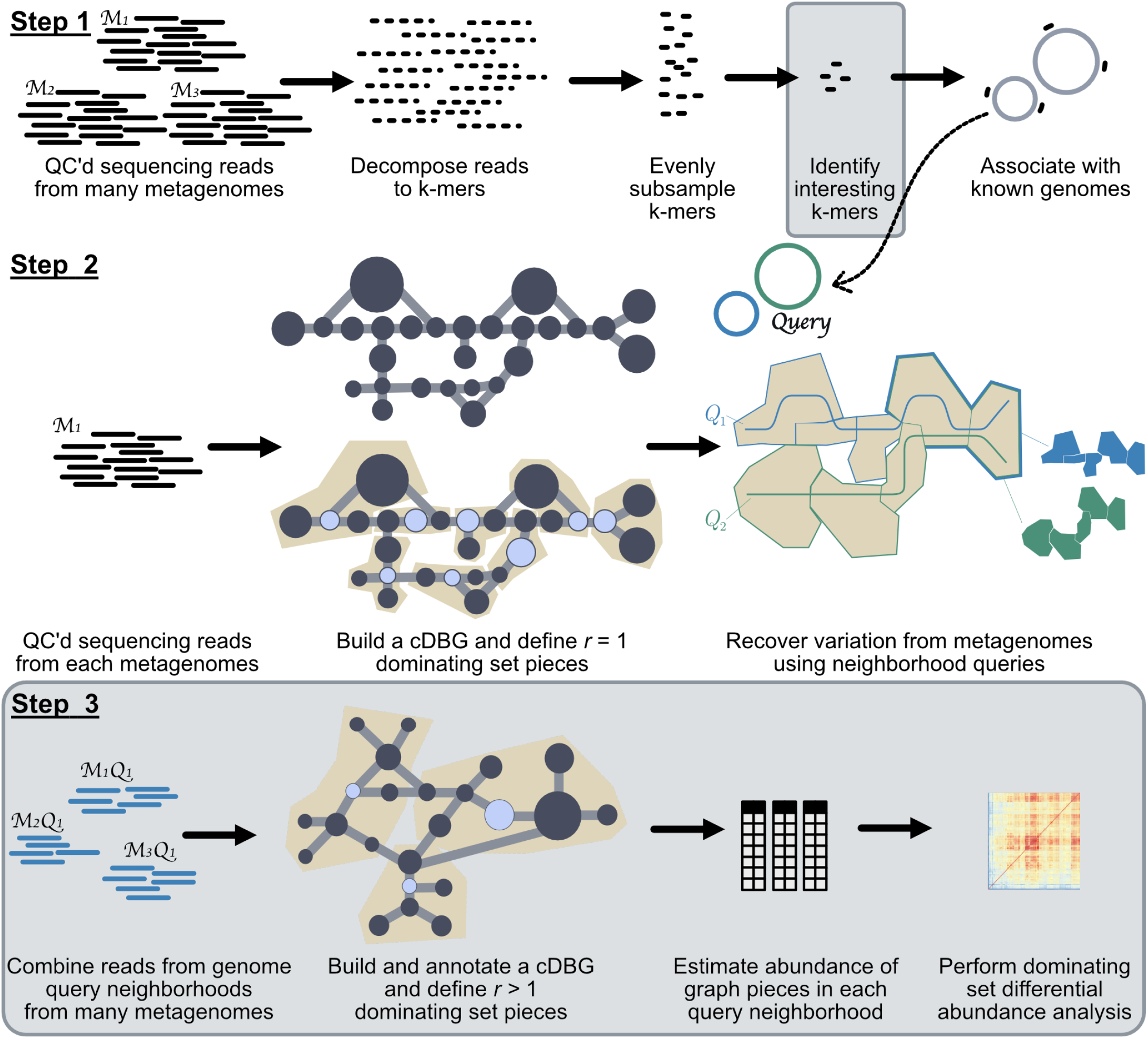
Overview of the metagenome analysis technique. Steps that are outlined in grey were developed in this paper. **Step 1:** Using quality controlled sequencing reads from many metagenomes, we decomposed reads into k-mers and subsample these k-mers using FracMinHash, thereby selecting k-mers that evenly represent the sequence diversity within a sample. We then identified interesting k-mers using random forests, and associated these k-mers with genomes in reference databases. **Step 2:** For each metagenome, we constructed a compact de Bruijn assembly graph (cDBG) that contains all k-mers from a metagenome. We used dominating sets to carve the graph into pieces. We queried this graph with genomes associated with interesting k-mers identified in Step 1, recovering sequence diversity nearby in the assembly graph. We refer to these sequences as genome query neighborhoods. Step 2 is the workflow published in [25]. **Step 3:** We combined genome query neighborhoods for a single genome from all metagenomes. We constructed a cDBG from these sequences, and used a dominating set with a large radius to carve the graph into large pieces. Here, we diagram construction of *r*=2 dominating set pieces, but in practice we used *r*=10. We estimated the abundance of k-mers in each metagenome for each dominating set piece, and used these abundances to perform differential abundance analysis.

Next, we developed an approach to perform differential abundance analysis directly on assembly graphs in order to recover all sequences that were differentially abundant in each IBD subtype when compared to nonIBD. Using the genomes identified by our k-mer association analysis, we first performed assembly graph genome queries to recover all sequences associated with a given species within a metagenome. For each genome query, we combined these sequences into a single assembly graph, which we refer to as a *metapangenome graph*. We estimated the abundance of each piece in this graph within each metagenome, and used these abundances to perform differential abundance analysis.

We applied this approach to the analysis of IBD gut microbiomes via meta-analysis. Meta-analyses have recently shown success in improving the power to detect microbial signatures of colorectal cancer [32,33,34]. To this end, we identified studies that performed metagenomic sequencing of individuals with CD, UC, or nonIBD and combined these to perform a meta-analysis (**Table 1**, **Table S1**). All studies profiled fecal gut microbiomes via Illumina shotgun metagenome sequencing. Individuals were from five distinct countries and seven cohorts (**Table 1**). In many studies, samples were taken in time series to profile disease progression or individual response to treatment. In these cases we included only the first sample in the time series so organized interventions would not skew our results. In addition, many of the nonIBD samples, particularly those from the iHMP, profiled sick individuals that were not diagnosed with IBD, meaning some of these samples are not healthy controls.

**Table 1:** Six IBD shotgun metagenome sequencing cohorts used in this meta-cohort analysis.

| Cohort | Name | Country | Total | CD | UC | nonIBD | Reference |
| --- | --- | --- | --- | --- | --- | --- | --- |
| iHMP | IBDMDB | USA | 106 | 50 | 30 | 26 | [5] |
| PRJEB2054 | MetaHIT | Denmark, Spain | 124 | 4 | 21 | 99 | [10] |
| SRP057027 | NA | Canada, USA | 112 | 87 | 0 | 25 | [8] |
| PRJNA385949 | PRISM, STINKi | USA | 17 | 9 | 5 | 3 | [12] |
| PRJNA400072 | PRISM, LLDeep, and NLIBD | USA, Netherlands | 218 | 87 | 76 | 55 | [30] |
| PRJNA237362 | RISK | North America | 28 | 23 | 0 | 5 | [31] |
| Total |  |  | 605 | 260 | 132 | 213 |  |

### K-mers are weakly predictive of IBD subtype

We first sought an approach to compare many metagenomes without relying on reference databases, *de novo* assembly, or annotations. We reasoned that FracMinHash sketches randomly subsample k-mers to allow comparisons, which may provide an unbiased approach to quickly compare across many metagenomes. In total, we profiled 7,376,151 subsampled k-mers across all samples in all cohorts, representing approximately 14 billion distinct k-mers in the original samples.

We detected variation correlated with IBD diagnosis in k-mer profiles of gut metagenomes from different cohorts. We calculated a pairwise distance matrix using angular distance between k-mer abundance profiles to assess sample diversity. We performed principle coordinate analysis and PERMANOVA with this distance matrix (**Figure 2 A, B**), using the variables study accession, diagnosis, library size, and number of k-mers observed in a sample (**Figure 2 B**). Study accounted for highest variation, emphasizing that technical artifacts or cohort diversity can introduce strong signals that may influence heterogeneity in results across IBD microbiome studies but that can be mitigated through meta-analysis [32]. The number of k-mers observed in a sample accounted for the second highest variation, possibly reflecting reduced diversity in stool metagenomes of CD and UC patients (reviewed in [35]). Diagnosis accounted for a substantial amount of variation as well, indicating that there is a small but detectable signal of IBD subtype in stool metagenomes.

To evaluate whether the variation captured by diagnosis is predictive of IBD subtype, we built random forest classifiers to predict CD, UC, or nonIBD subtype. To assess whether disease signatures generalize across study populations, we used a leave-one-study-out cross-validation approach where we built and optimized a classifier using five cohorts and validated on the sixth. We built each model six times to hone in on cross-study and cross-model signal. Given the high-dimensional structure of this data set (e.g. many more k-mers than metagenomes), we first used the vita method of variable selection to narrow the set of predictive k-mers in the training set [36,37]. Variable selection reduced the number of k-mers used in each model by two orders of magnitude, from 7,376,151 to 28,684-41,701 (mean = 35,673.1, sd = 4090.3) (**Figure 2 C**).

Using this reduced set of k-mers, we optimized each random forests classifier on the training set, producing 36 optimized models. We validated each model on the left-out study. The accuracy on the validation studies ranged from 49%-77% (**Figure 2 D**), outperforming a previously published model built on metagenomic data alone [30].

**Figure 2:**
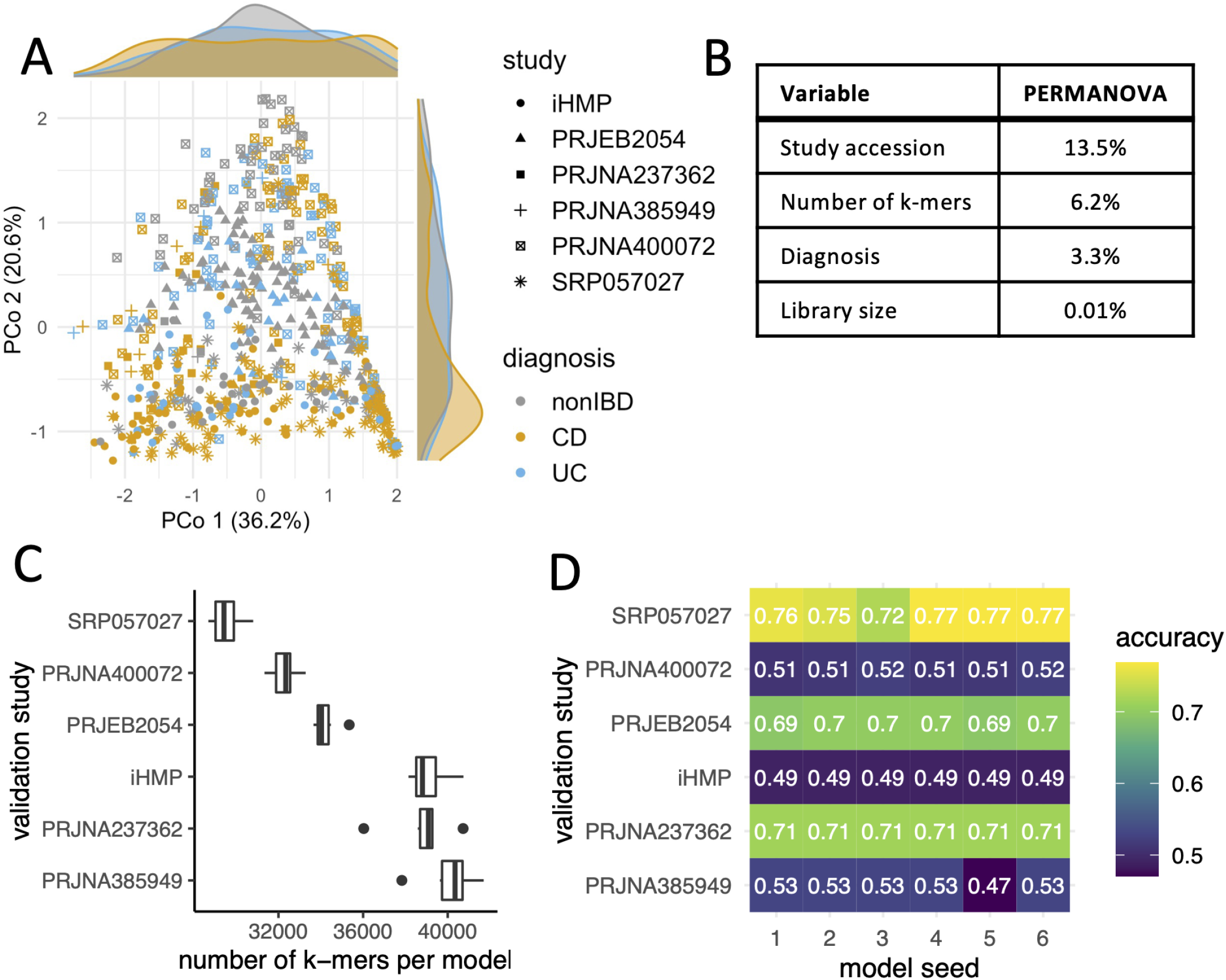
Long nucleotide k-mers retain information about IBD subtype classification. **A.** Principal coordinate analysis of distance matrices obtained from comparing FracMinHash sketches with abundances and **B.** PERMANOVA results that explain the variance in the principal coordinate analysis. Number of k-mers refers to the number of k-mers in a sketch, while library size refers to the number of raw reads per sample. All tests were significant at p < .001. **C.** Box plots indicating the number of k-mers used to build each random forests model. Variable selection using the vita method reduced the number of k-mers used to build each model. **D.**Heatmap indicating accuracy of each model on the left-out validation study. Model performance varied by validation study, but models predicted IBD subtype better than chance (1/3).

To understand which genomes were responsible for disease signatures detected by our models, we anchored k-mers in the models against genomes in reference databases using sourmash gather [21]. Sourmash gather determines the minimum set of genomes in a database necessary to cover all of the k-mers in a query [21]. We used the GTDB rs202 representatives database, which contains bacterial and archaeal genomes, and the GenBank viral, fungal, and protozoa databases. We found that a substantial fraction of genomes were shared between models, indicating there is a consistent biological signal captured among classifiers: of 3,889 total genomes detected across all classifiers, 360 genomes were shared between all classifiers (**Figure 3**, **Figure S1**,**Table S2**). The presence of shared k-mers between classifiers indicates that there is a consistent biological signal in metagenomes for IBD subtype between cohorts.

K-mers that anchored to these shared genomes represented 65% of all k-mers used to build the optimized classifiers, but accounted for an outsize proportion of variable importance in the optimized classifiers. After normalizing variable importance across classifiers, 74% of the total variable importance was held by these k-mers. These k-mers contribute a large fraction of predictive power for classification of IBD subtype, and the genomes in which they are found represent a microbial core that contains predictive power in IBD subtype classification.

Given that 360 genomes anchored the majority of k-mers and variable importance across all models, we were curious whether a smaller number of genomes could still retain the majority of variable importance. Limiting genomes to those that could hold at least 1% of the normalized variable importance, we found that 54 genomes accounted for 50% of the variable importance (**Figure 3**, **Figure S1**, **Figure S2**, **Table S2**). We assumed these genomes represent the strongest candidates for discriminating IBD subtype and focused on them for the remainder of our analyses.

**Figure 3:**
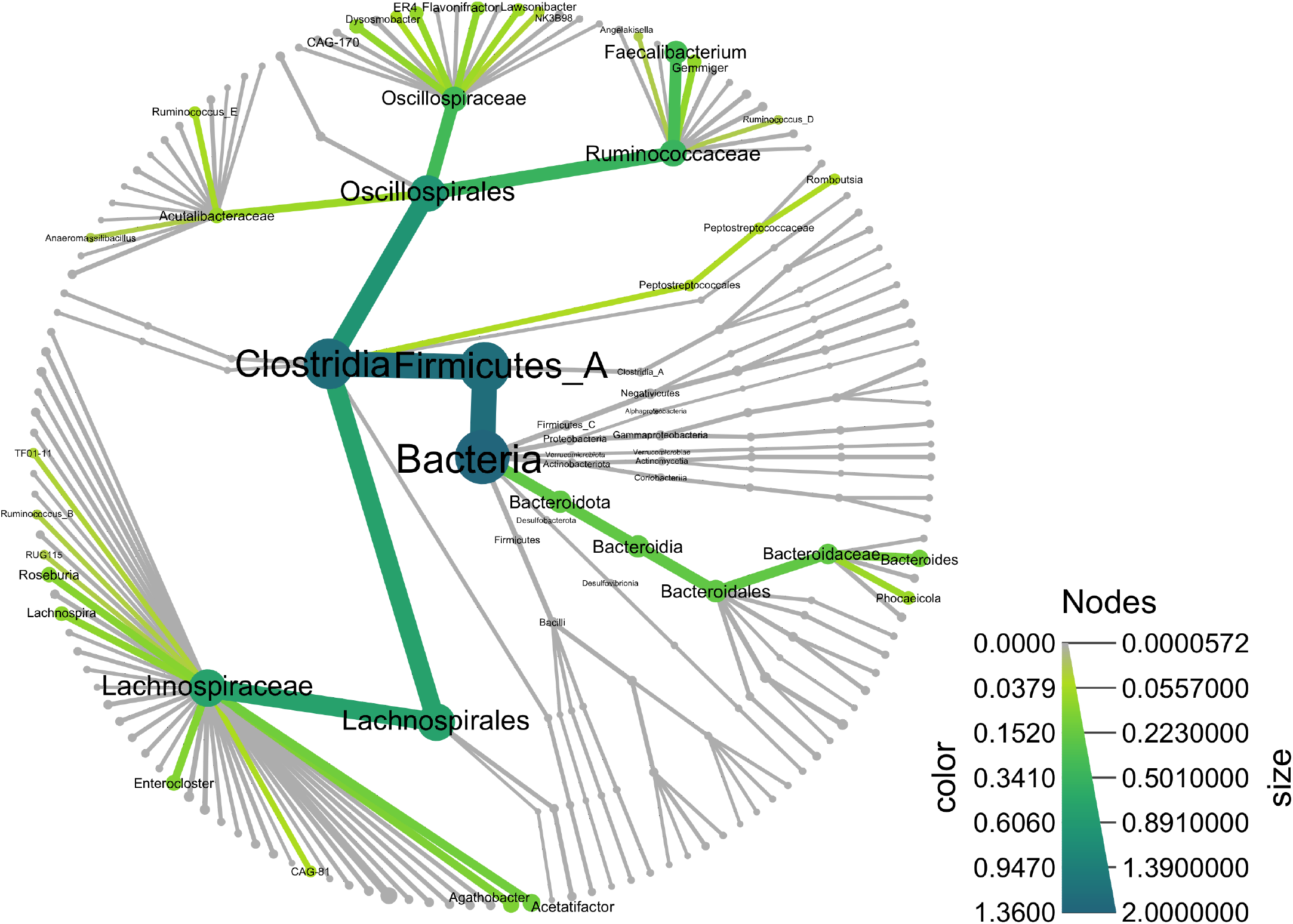
Tree of bacterial species that were predictive of IBD subtype in all models. Nodes are summarized to the genus level. All taxa up to the class level are labelled. Taxa that could account for at least 1% of the normalized variable importance across random forests models are colored and labelled. Node size and node color reflect potential normalized variable importance attributable to each taxonomic lineage with larger node sizes and darker color representing larger variable importance; while normalized variable importance across models sums to one, some sequences are shared across genomes making the total potential variable importance across all genomes larger than one.

### Genome queries into metagenome assembly graphs recover neighborhoods of sequence variation and establish species umbrellas

While we were able to identify the majority of k-mers that were important for predicting IBD subtype, 26% of k-mers remained unannotated. We hypothesized that these k-mers represented strain variable sequences not in reference databases but belonging to species represented by annotated k-mers. To investigate this hypothesis, we performed genome queries on assembly graphs of each metagenome using the 54 candidate genomes that discriminated IBD subtype (**Figure 1**). Assembly graph genome queries recover sequences in a metagenome that match the query, as well as those that are nearby in the assembly graph, making queries akin to but more general than read mapping against reference genomes (**Figure 1**) [25]. The resulting genome query neighborhood represents a species-level umbrella that contains sequence variation from the metagenome associated with a query.

After performing genome queries, we re-anchored k-mers against the resulting query neighborhoods as well as the databases used previously. We observed that the fraction of unassigned k-mers decreased from 26% to 8% (**Figure 4**), supporting our hypothesis that many of these k-mers are sequence variants belonging to species identified in k-mers important for predicting IBD subtype. We further observed that many other k-mers previously anchored by other genomes were reassigned to the genome query neighborhoods (**Figure 4**). This suggests that the genome queries create species umbrellas that represent sequence variation for the query genome itself as well as other closely related genomes that occur within a metagenome.

**Figure 4:**
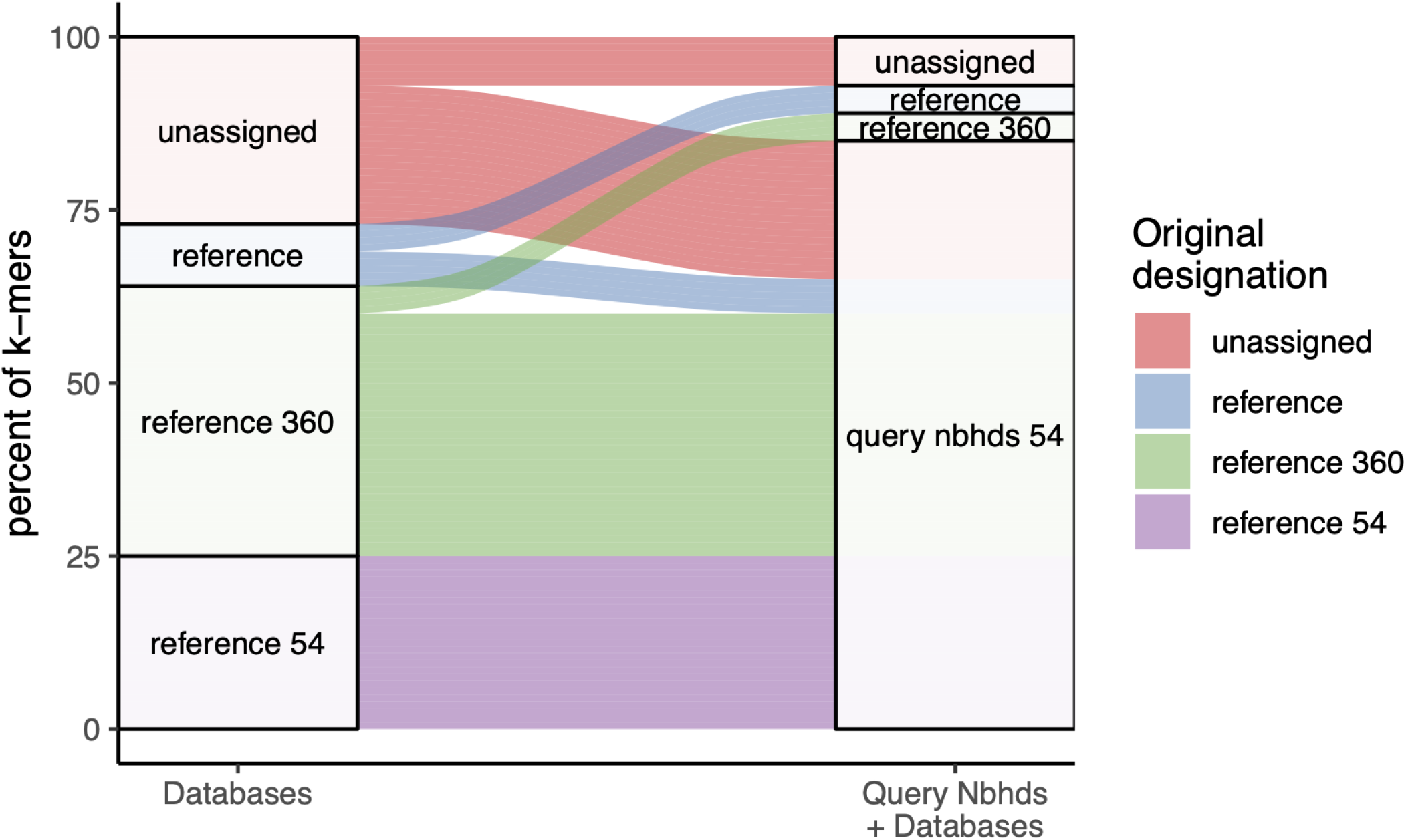
Alluvial plot depicting the set of genomes that anchored k-mers that were important for predicting IBD subtype. The blocks on the left represent the breakdown of k-mer assignations from greedy exact matching against databases alone, while the blocks on the right represent k-mer assignations after metagenome assembly graph queries.

### IBD gut microbiomes have decreased diversity in strict anaerobes that is punctuated by strain switches for some facultative anaerobes

After recovering all sequences in metagenomes in the neighborhoods of the species that discriminate IBD subtypes, we next sought to determine the specific genome segments that were differentially abundant in IBD. Differential abundance analysis is a common step in metagenome analysis, however it is typically applied to gene counts [38,39], which requires assembly or mapping prior to abundance estimation. To avoid assembly or mapping and the accompanying loss of reads [40], we developed an abundance estimation approach that works directly on the assembly graphs, enabling differential abundance analysis from the assembly graph. Our abundance estimation approach was based on *r*-dominating sets, an algorithm introduced in [25] that efficiently computes the dominating nodes in a cDBG so that every node is no more that distance *r* from a dominator. The dominating set is used to carve the graph into pieces, each of which contains one dominating node. Here, we first build a species-level assembly graph that contains neighborhood sequences for a given genome across all metagenomes, which we call a *metapangenome graph*. We then partition the graph into pieces using a large radius (*r* = 10). The large radius carves the graph into pieces that average 103 k-mers in size. We next estimated the abundance of each piece within each metagenome using average k-mer abundance. We also annotated the graph pieces using using k-mer overlap between genes of known function and graph pieces. Using this information, we performed dominating set differential abundance analysis using corncob [41], a statistical package that tests for differential relative abundance in the presence of variable sequencing depth and excessive zeroes for unobserved observations, conditions which occur in abundances from dominating sets. We tested differential abundance at the 5% significance level after correcting for multiple testing (see methods).

We applied this method for each of our genome queries, building 54 metapangenome graphs and performing dominating set differential abundance analysis on each. We tested for differential abundance in pieces that occurred in at least 100 metagenomes, since we sought differences that characterized the majority of our samples within a group. Note that corncob fits a model for each dominating set piece and therefore does not require abundance information for all pieces [41]. On average, this condition was met in 6.4% of dominating set pieces. Focusing on pieces that occurred in many metagenomes increased the average piece size to 1088 k-mers, which is similar to the average bacterial gene length of approximately 1000 base pairs [42] and should enable biologically meaningful comparisons across groups.

**Figure 5:**
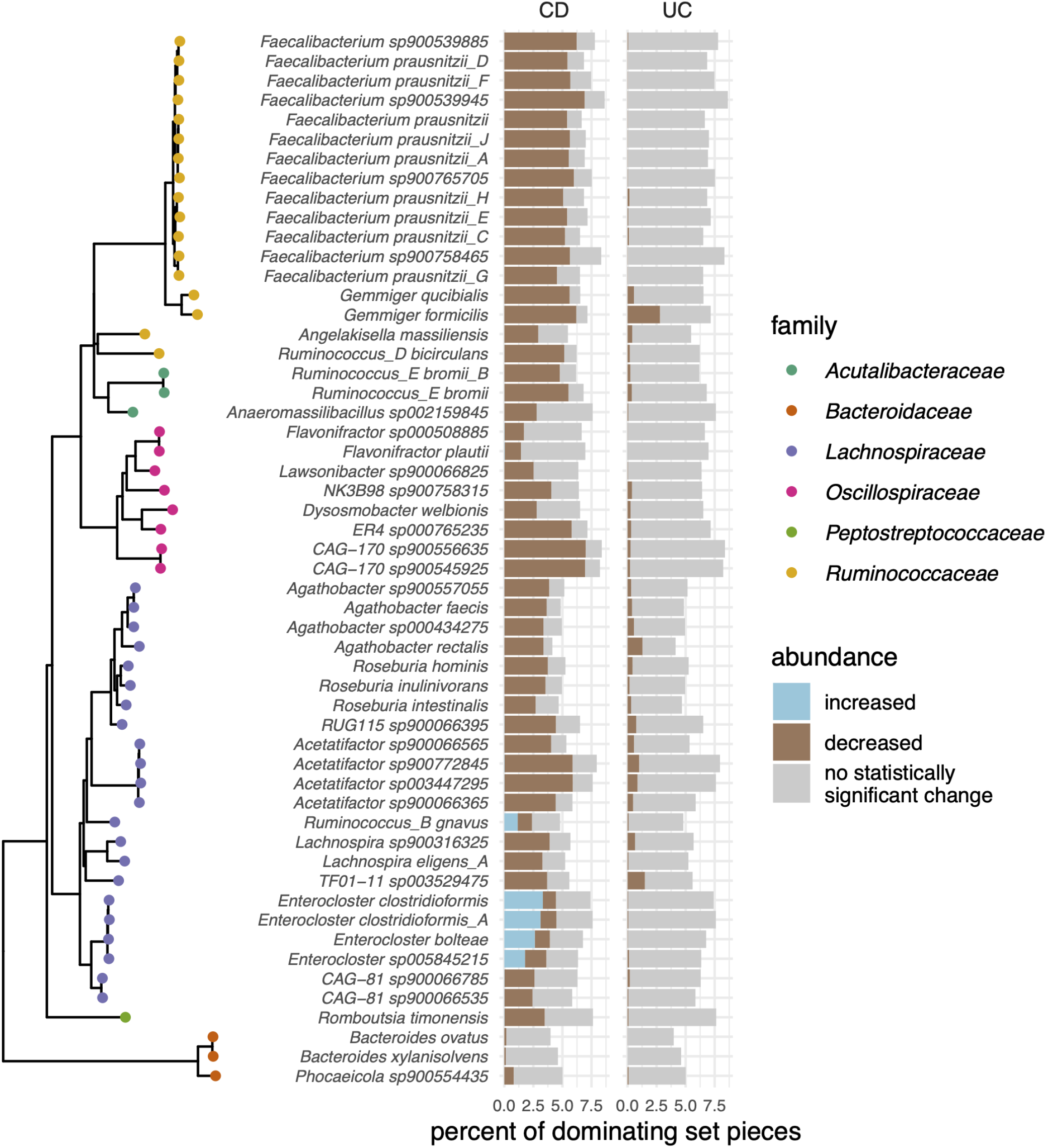
Dominating set differential abundance analysis revealed genome segments that were significantly different in CD and UC compared to nonIBD. Results are organized by GTDB taxonomy, with a tree representing the 54 species and colored by family on the far left. The percent of dominating set pieces tested is labelled in grey, and the percent of significantly differentially abundant pieces are colored by increased (blue) or decreased (brown) abundance.

We found that the majority of species decreased in abundance in CD, and to a lesser extent, UC (**Figure 5**). Many of these species are generally regarded as beneficial bacteria. For example, nine of the 54 genomes we investigated were *Faecalibacterium prausnitzii*, the phylogroups of which are separated in the GTDB taxonomy but combined into a single species in the NCBI taxonomy. *F. prausnitzii* is a key butyrate producer in the gut and plays a crucial role in reducing intestinal inflammation [43]. Similarly, *Acetatifactor* is a bile-acid producing bacteria associated with a healthy gut, but limited evidence has associated it with decreased abundance in IBD [44]. These species, as well as others that decreased in abundance in IBD, are strictly anaerobic (**Figure 5**), so these observed trends are consistent with a shift toward oxidative stress during disease that is intolerable for many gut microbes [45].

A substantial fraction of dominating set pieces were more abundant in CD than nonIBD in five metapangenome graphs (**Figure 5**). These graphs represented sequences from species *R. gnavus, Enterocloster bolteae, Enterocloster sp005845215, Enterocloster clostridioformis*, and *Enterocloster clostridioformis_A*. We posit that the increased abundance in some genomic segments amid the decrease in abundance of others represents strain switching that occurs in CD.

In support of this, when we annotated the differentially abundant pieces using KEGG orthologs, we found that in some cases pieces that were increased in abundance and pieces that were decreased in abundance were annotated with the same ortholog (mean = 1453.8, sd = 727.2 pieces representing mean = 64, sd = 23.8 orthologs per graph, **Figure S3, Table S3**). These genes likely represent the portion of the core genome shared by the strain(s) that is more abundant in CD and the strain(s) that is more abundant in nonIBD, but that is encoded by distinct sequences. Some shared annotations encoded single copy marker genes [46]. To confirm that multiple strains of the same species were represented by these sequences, we queried with these marker gene sequences, extracted the reads associated with those graph pieces, and mapped the reads back to the marker gene sequence. We then selected reads that aligned to the same portion of the marker gene and contained single nucleotide variants, and BLASTed those reads against the NCBI nr database. For the subset of reads that we tested, we confirmed that different strains of the same species were the best matches. This demonstrates that mulitple strains of the same species were present in each species graph, and that these strains were differentially abundant in CD compared to nonIBD.

In contrast to the annotations that were identified among sequences that were increased and decreased in abundance, many orthologs were only annotated among the pieces that were increased in abundance in CD (mean = 1193.4, SD = 155.7, **Table S4**). Many of the same pathways were enriched among these orthologs across the five species, including those associated with oxidative stress response (cysteine and methionine metabolism, the pentose phosphate pathway) (**Figure S3**). The oxidative branch of the pentose phosphate pathway regenerates NADPH, while cysteine is a precursor for the antioxidant glutathione. Given this, we investigated the presence of reactive oxygen species-scavenging orthologs [47]; sequences encoding superoxide dismutase (K04565), thioredoxin reductase (K00384), and peroxiredoxin (K03386) were increased in abundance in CD for all five species. Additionally, many enriched pathways were associated with virulence (quorum sensing, flagellar assembly, bacterial chemotaxis, vancomycin resistance). Enrichment of specific metabolic pathways is consistent with functional specialization of strains in different environmental niches [48].

While dominating set differential abundance analysis identified genomic sequences that were more abundant in CD, the nature of short shotgun metagenomic sequencing reads precludes haplotype phasing or lineage resolution [15], meaning our results likely represent genomic variants from many distinct genomes that would not all naturally occur together in a single isolate genome. Therefore, to identify isolate genomes that contain the genomic sequences that were more abundant in CD, we searched the GTDB rs202 database with the significantly differentially abundant sequences. On average, the top matching isolate genome contained 63% of the sequences that were more abundant in CD (**Table 2**).

**Table 2:**
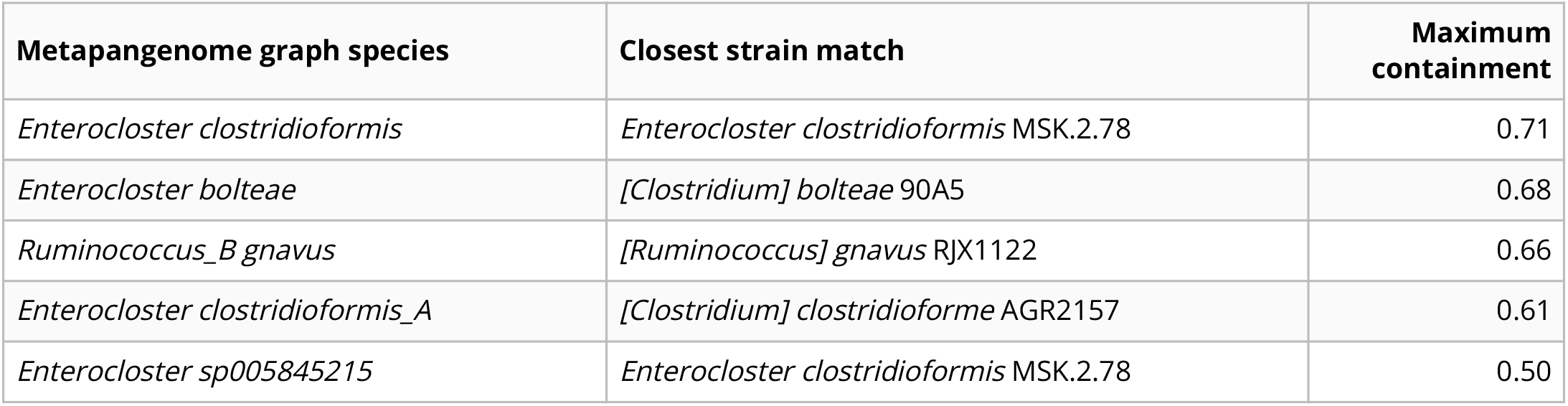
Maximum containment between sequences that were increased in abundance in CD and isolate genomes.

One aerotolerant clade of *R. gnavus* was previously identified as being enriched in CD [12], and has been shown to produce a polysaccharide that induces the inflammatory cytokine TNF-alpha [13]. The three isolate genomes we identified as containing the highest amount of sequences that were increased in abundance in CD were among those that have been shown to induce TNF-alpha secretion (RJX1122, RJX1127, RJX1128) [14]. This suggests our method identified the same strain switch previously discovered to occur in IBD [12,13,14]. In further support of this, we found that 17 of the 23 genes in the operon that encodes the proteins responsible for producing the inflammatory polysaccharide were annotated in the dominating set pieces that were more abundant in CD. These genes were encoded across 66 dominating set pieces, with multiple neighboring genes in the operon annotated in 6 of these dominating set pieces.

For two of the four *Enterocloster* species, the top matching isolate sequence was the same (*Enterocloster clostridioformis* MSK.2.78). This points to overlap in the genomic sequences we identified as differentially abundant across these metapangenome graphs. Indeed, the average Jaccard similarity between the sequences that were increased in CD in the *Enterocloster* graphs was 0.53, while the average max containment was 0.74. Given that a Jaccard similarity of 0.1 is required to recover at least 80% of a genome via assembly graph query, which approximately corresponds to an average nucleotide identity of 93% [49], and that species boundaries in GTDB are drawn at 95% average nucleotide identity [50], the metapangenome graphs likely store genomic sequences associated with both the query genome species and closely related species. However, the metapangenome graphs presented here, as well as the differentially abundant sequences, contain both common and distinct nucleotide sequences, suggesting that multiple closely related *Enterocloster* species/genomes are associated with CD. Taken together, our ability to recover a previously validated sub-species association with IBD (*R. gnavus*) suggests that the three new *Enterocloster* isolates we identified should be further investigated for their potential role in eliciting CD-like symptoms in the gut.

### Genomic sequences that are differentially abundant in IBD are not exclusive to IBD

Since genome sequences belonging to many species were differentially abundant from nonIBD in CD and UC, we next investigated whether there was a disease-specific microbiome in CD or UC – i.e., whether there are sequences from a species that were only observed in IBD. Using FracMinHash sketches from the differentially abundant sequences, we identified the differentially abundant sequences in each metagenome and compared their occurrence and distribution across diagnoses.

In general, we found no evidence for disease-specific sequences among the 54 species we investigated. Using FracMinHash sketches of the differentially abundant sequences for each species, we counted the number of k-mers that were observed in different sets of diagnoses. We observed almost all sequences in at least some CD, UC, and nonIBD metagenomes (**Figure 6**, **Figure S4**). Across all species, an average of 14.9% differentially abundant k-mers were unobserved in either CD, UC, or nonIBD. These results in part explain heterogeneous study findings in previous IBD gut microbiome investigations.

**Figure 6:**
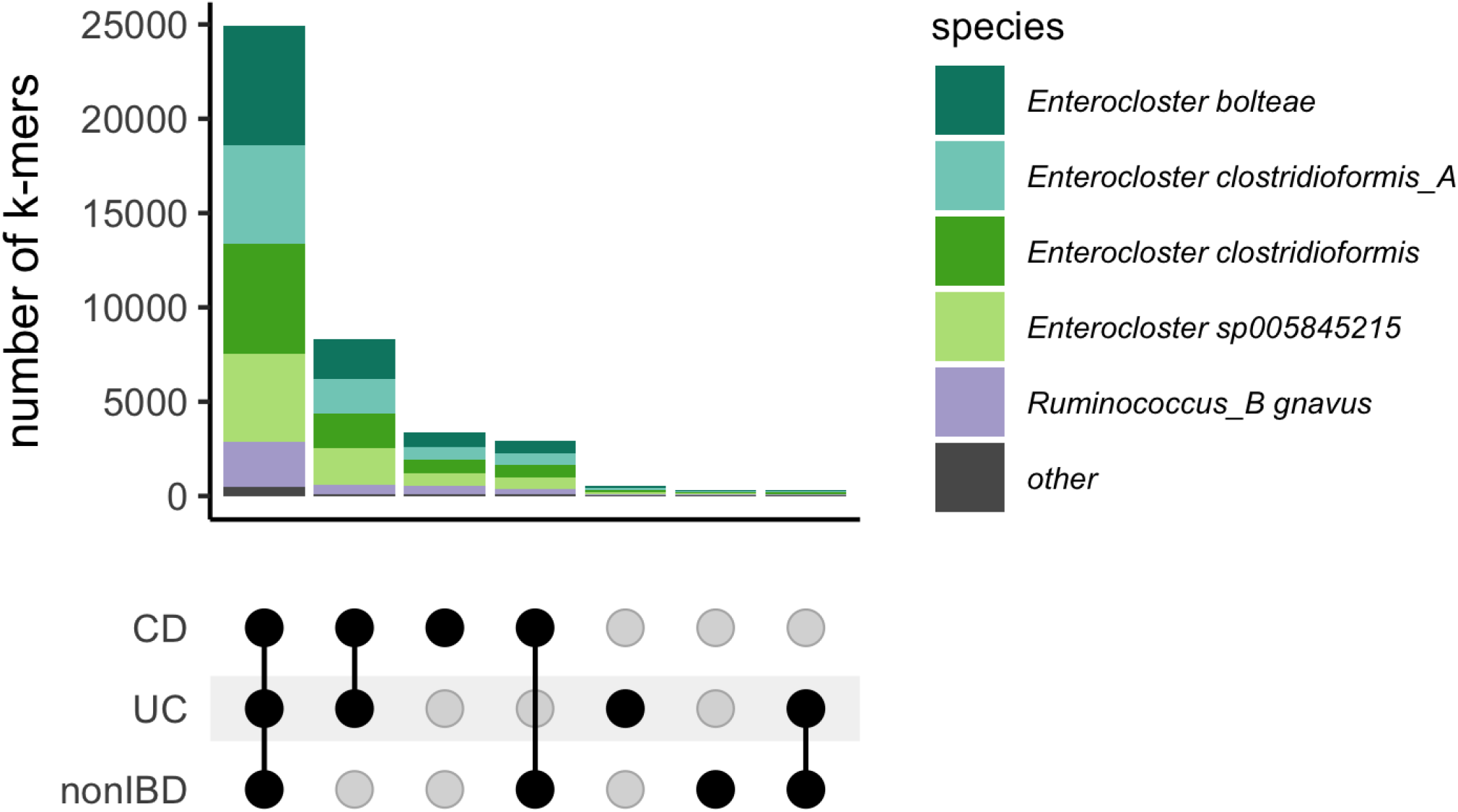
Most differentially abundant sequences occur in metagenomes of individuals diagnosed with CD, UC and non-IBD. Upset plot of k-mers that were increased in abundance in CD and their occurrence in CD, UC, and nonIBD metagenomes. The bottom half of the plot highlights which diagnoses are included in each set, while the bar chart in the top half of the plot shows the number of k-mers that were observed in that set. The bar chart is colored by the metapangenome species graph in which the sequence was differentially abundant.

## Discussion

In this paper, we present an assembly-free metagenome analysis framework for group association discovery that is minimally reliant on reference databases. Our approach uses k-mers to discover genomes associated with groups of metagenomes, and then recovers sequence variation from those genomes and closely related genomes in the metagenomes. These sequences are organized in a “metapangenome graph” which is then used to perform differential abundance analysis to discover specific genomic sequences that differ between groups.

We applied this method to perform a meta-analysis of fecal gut microbiome metagenomes from individuals with CD, UC, and nonIBD and uncovered cross-study microbial signatures of IBD subtype. The underlying etiology of IBD remains poorly understood with inconsistent microbiological findings produced from different studies [4]. The signatures we identified demonstrate consistent loss of diversity of specific microorganisms, particularly in CD. Among the background of generalized loss of diversity, we observed that some genomic sequences increased in abundance while others decreased in abundance for five species in CD. This pattern is consistent with strain switching, where one strain is more abundant in CD and another is more abundant in nonIBD. For one species we identified, *R. gnavus*, this strain switching behavior was previously discovered via isolate sequencing and metagenome mapping [12]. Our recovery of this pattern demonstrates the utility of our approach for discovering sub-species level associations from metagenomic sequences alone, and opens the door for additional discovery. Indeed, we identified four additional species where strain switching occurred in CD.

While our approach identified genomic sequences that were more abundant in CD than nonIBD, the nature of short read sequences precludes haplotype or lineage resolution directly from the metagenomic data analyzed here. To circumvent this issue, we identified isolate genomes that encoded all of the genomic sequences that were more abundant in CD. These isolate genomes represent candidate organisms for further research into the microbial component of CD pathophysiology. As high fidelity long read sequencing of microbiomes becomes increasingly common [15], long reads can be integrated into the approaches introduced here, enabling lineage-resolved association detection directly from sequencing data.

While we found conserved signatures in IBD subtype, we found no evidence for disease-specific sequences within the gut microbiome. The observation that almost all differentially abundant sequences for a given species occur in CD, UC, and nonIBD suggests the presence of ecotypes - subspecies that are adapted to different environments – rather than pathotypes - subspecies associated with a specific disease. These patterns in part explain the inconsistent results generated in IBD subtype characterization, where no consistent microbiological signal has emerged in human gut microbiomes other than loss of diversity [4].

Our models consistently performed the most poorly on the iHMP cohort. The iHMP tracked the emergence and diagnosis of IBD through time series profiling of emergent cases [5]. We selected the first sample in each time series for this analysis, and given the relatively poor performance of these models, this may suggest that disease onset is a distinct biological process. However, the inclusion of the iHMP cohort in this analysis ensured that not all nonIBD samples were healthy controls and some fraction were symptomatic cases that did not result in an IBD diagnosis [5].

While we apply our pipeline to IBD classification, it is extensible to other large meta cohorts of metagenomic sequencing data. This method may be particularly suitable for diseases such as colorectal cancer, where a recent meta-analysis using a marker gene approach was successful in classifying colorectal samples from healthy controls [32]. Beyond classification of disease, our method may bring strain-level resolution and generate hypotheses for further research. This may be particularly useful in the context of tumor microbiomes as previous research has demonstrated that strain-specific *Helicobacter pylori* and human papillomaviruses are risk factors for or directly transmit certain cancers [51]. Strain-resolved methods may further this area of research.

The methods we used to perform the k-mer association analysis are modular and may be improved by substituting parts of the pipeline with different approaches. For example, we used abundances from long nucleotide k-mers (*k* = 31) – which capture species-level sequence similarity [20] – as our features. K-mers constructed from protein or other reduced alphabets may improve accuracy, as we would expect more shared sequence content between metagenomes as well as a better representation of functional content [52]. While this may improve classification accuracy, switching to reduced alphabet k-mers may not be desirable in the context of strain-specific differences which may be obscured by these degenerate representations. Similarly, while we used random forests to to perform k-mer association analysis, other machine learning or statistical techniques may improve classification accuracy. These approaches remain areas of future research.

The first part of the pipeline is disconnectable from the second part of the pipeline - that is, the discovery of discriminatory genomes between groups is not a prerequisite for dominating set differential abundance analysis as query genomes could be selected arbitrarily. Therefore, the assembly graph differential abundance approach presented here could be applied to metagenomes for samples originating from diverse environments. The requirements for the application of dominating set differential abundance are threefold. First, there must be sufficient samples for comparison (e.g., a minimum of three cases and three controls, with the typical caveats for small sample sizes [53]). Second, we must have a genome with which to query the graph. And third, we must have sufficient compute resources to run spacegraphcats [25]. These requirements make the application of dominating set differential abundance analysis available to metagenomes from diverse environments, not just the well-studied human gut microbiome.

While we present an initial pipeline that enables differential abundance analysis directly on assembly graphs, we identified several areas where our approach could be improved. First, implementing approaches to better control graph piece size would be beneficial. We heuristically selected *r* = 10 to build the metapangenome graphs because this radius produced dominating set pieces with approximately the same number of k-mers as the average bacterial gene for pieces that were present in many metagenomes. However, the radius needed to meet this condition may change depending on the diversity of the sequencing reads used to build the metapangenome graph. Diversity increases with the complexity of the sequenced community, sequencing depth, and the number of communities observed. Algorithms that either automatically select a radius that achieves a user-specified average piece size, or that produce more consistent piece sizes independent of diversity of the sequencing data would provide finer control of the graph structure and subsequent sequence comparisons. Further, switching from a cDBG to a de Bruijn graph as the base spacegraphcats graph structure could lead to more consistent piece sizes; cDBGs have variable node sizes because they combine nodes without branching paths, while every node in a de Bruijn graph contains one k-mer. Second, improving RAM efficiency at high radii would enable more diversity to be represented in individual graphs. In order to build the metapangenome graphs, we first hard-trimmed the input sequences to remove low abundance k-mers, thereby decreasing the RAM needed to construct each graph. Algorithmic changes that improve RAM efficiency at high radii would obviate the need for hard trimming, and increase the amount of diversity that could be represented in a single graph. Similarly, improved performance would allow dominating set differential abundance analysis to be performed directly on groups of metagenomes without the need to first identify species of interest via genome queries.

## Methods

All code associated with our analyses is available at github.com/dib-lab/2020-ibd (DOI:10.5281 /zenodo.6783208). An example repository for dominating set differential abundance analysis is available at github.com/dib-lab/2022-dominating-set-differential-abundance-example (DOI:10.5281 /zenodo.6783363).

### IBD metagenome data acquisition and processing

We searched the NCBI Sequence Read Archive and BioProject databases for shotgun metagenome studies that sequenced fecal samples from humans with Crohn’s disease, ulcerative colitis, and healthy controls. We included studies sequenced on Illumina platforms with paired-end chemistries and with sample libraries that contained greater than one million reads. For time series intervention cohorts, we selected the first time point to ensure all metagenomes came from treatment-naive subjects.

We downloaded metagenomic FASTQ files from the European Nucleotide Archive using the “fastq_ftp” link and concatenated fast files annotated as the same library into single files. We also downloaded iHMP samples from idbmdb.org. We used Trimmomatic (version 0.39) to adapter trim reads using all default Trimmomatic paired-end adapter sequences (`ILLUMINACLIP: {inputs/adapters. fa}:2:0:15`) and lightly quality-trimmed the reads (`MINLEN: 31 LEADING: 2 TRAILING:2 SLIDINGWINDOW:4:2`)[54]. We then removed human DNA using BBMap and a masked version of hg19 [55]. Next, we trimmed low-abundance k-mers from sequences with high coverage using khmer’s `trim-low-abund.py` [56].

Using these trimmed reads, we generated FracMinHash signatures for each library using sourmash (k-size 31, scaled 2000, abundance tracking on) [57]. FracMinHash sketching produces compressed representations of k-mers in a metagenome while retaining the sequence diversity in a sample [21,23]. This approach creates a consistent set of k-mers across samples by retaining the same k-mers when the same k-mers were observed. This enables comparisons between metagenomes. We refer to FracMinHash sketches as *signatures*, and to each sub-sampled k-mer in a signature as a *k-mer*. At a scaled value of 2000, an average of one k-mer will be detected in each 2000 base pair window, and 99.8% of 10,000 base pair windows will have at least one k-mer representative. We selected a k-mer size of 31 because of its species-level specificity [20]. We retained all k-mers that were present in multiple samples.

### Principle Coordinate Analysis

We used Jaccard distance and angular similarity as implemented in `sourmash compare` to pairwise compare FracMinHash signatures. We then used the `dist()` function in base R to compute distance matrices. We used the `cmdscale()` function to perform principle coordinate analysis [58]. We used ggplot2 and ggMarginal to visualize the principle coordinate analysis [59]. To test for sources of variation in these distance matrices, we performed PERMANOVA using the `adonis` function in the R vegan package [60]. The PERMANOVA was modeled as `~ diagnosis + study accession + library size + number of k-mers`.

### Random forests classifiers

We built random forests classifiers to predict CD, UC, and non-IBD status using FracMinHash signatures. We transformed sourmash signatures into a k-mer (hash) abundance table where each metagenome was a sample, each k-mer was a feature, and abundances were recorded for each k-mer for each sample. We normalized abundances by dividing by the total number of k-mers in each FracMinHash signature. We then used a leave-one-study-out validation approach where we trained six models, each of which was trained on five studies and validated on the sixth. We built each model six times, each time using a different random seed. To build each model, we first performed vita variable selection on the training set as implemented in the Pomona and ranger packages [37,61]. Vita variable selection reduces the number of variables (e.g. k-mers) to a smaller set of predictive variables through selection of variables with high cross-validated permutation variable importance [36]. It is based on permutation of variable importance, where p-values for variable importance are calculated against a null distribution that is built from variables that are estimated as non-important [36]. This approach retains important variables that are correlated [36,62], which is desirable in omics-settings where correlated features are often involved in a coordinated biological response, e.g. part of the same operon, pathways, or genome [63,64]. Using this smaller set of k-mers, we then built an optimized random forests model using tuneRanger [65]. We evaluated each validation set using the optimal model, and extracted variable importance measures for each k-mer for subsequent analysis. To make variable importance measures comparable across models, we normalized importance to 1 by dividing variable importance by the total number of k-mers in a model and the total number of models.

### Anchoring predictive k-mers to genomes

We used sourmash `gather` with parameters `k 31` and `-scaled 2000` to anchor predictive k-mers to known genomes [57]. Sourmash `gather` searches a database of known k-mers for matches with a query [21]. We used the sourmash GTDB rs202 representatives data base (https://osf.io/w4bcm/download). To calculate the cumulative variable importance attributable to a single genome, we used an iterative winner-takes-all approach. The genome with the largest fraction of predictive k-mers won the variable importance for all k-mers contained within its genome. These k-mers were then removed, and we repeated the process for the genome with the next largest fraction of predictive k-mers. To genomes that were predictive in all models, we took the union of predictive genomes from the 36 models. We filtered this set of genomes to contain only those genomes with a cumulative normalized variable importance greater than 1%.

### *r*-dominating sets

The original spacegraphcats publication defined the dominating set as a set of nodes in the compact de Bruijn graph (cDBG) such that every node is a distance-1 neighbor of a node in the dominating set [25]. However, the algorithms as implemented allow this distance to be flexible and tunable [25]. We refer to the largest distance that any node may be from a member of the dominating set as the *radius, r*. Increasing the radius increases the average piece size while reducing the total number of pieces in the graph.

### Genome neighborhood queries with spacegraphcats

To recover sequence variation associated with genomes that were correlated with IBD subtype, we used spacegraphcats `search` to retrieve k-mers in the compact de Bruijn graph neighborhood of each genomes (*r* = 1, *k* = 31) [25]. We then used spacegraphcats `extract_reads` to retrieve the reads and `extract_contigs` to retrieve unitigs in the cDBG that contained those k-mers, respectively.

### Construction of the metapangenome graph

After retrieving genome neighborhood sequences from each metagenome, we combined these sequences to build a single metapangenome graph (*r* = 10, *k* = 31). We increased the radius of the metapangenome graph to produce larger level 1 dominating set pieces and to overcome highly articulated cDBGs resulting from an abundance of sequencing data. While working with single-species metapangenome graphs from many metagenomes reduced the graph size compared working with complete metapangenome graphs, we performed two preprocessing steps prior to the metapangenome graph generation. We combined all genome query neighborhood reads and performed digital normalization and then truncated reads at k-mer that was not present in the data set at least 4 times. These are heuristic steps that we believe are unlikely to remove biologically important sequences.

### Annotating the metapangenome graph

We implemented an approach to annotate dominating set pieces in spacegraphcats assembly graph. This approach is implemented in spacegraphcats as `multifasta_queries`. This approach relies on k-mer overlap between sequences in a reference multifasta file and nodes in the cDBG and is executed in a two-step approach. First, `search. index_cdbg_by_multifasta. py` identifies all cDBG nodes that match to the k-mers in a FASTA sequence and promotes those annotations to all cDBG records in the dominating set piece. Then, `search/extract_cdbg_by_multifasta. py` extracts and summarizes information about these annotations and outputs it to a CSV fìle.

We applied this annotation approach to the metapangenome graphs for species that were more abundant in CD. To generate a reference multifasta gene fìle to transfer annotations from, we first downloaded all genomes of the species represented in the metapangenome graph and in the GTDB rs202 database. We annotated open reading frames (ORFs) in these genomes using bakta [66], combined and clustered predicted ORFs using cdhit-est [67], and performed ortholog annotation using eggnog [68].

### Calculating abundances metagenome abundances of dominating set nodes in the metapangenome graph

We implemented an approach to calculate k-mer abundances for each graph piece in the level 1 dominating set. This approach is implemented in spacegraphcats in `search/count_dominator_abundance.py`. Using a spacegraphcats assembly graph and a set of reads from a metagenome, the abundance of each dominating set piece is calculated by summing the abundance of every k-mer in that piece within the metagenome.

We applied this abundance estimation approach to each metapangenome graph, estimating the abundance of each dominating set piece within each metagenome.

### Performing dominating set differential abundance analysis

We used Corncob to perform dominating set differential abundance analysis [41]. Corncob tests for differential relative abundance in the presence of variable sequencing depth and excessive zeroes for unobserved observations, conditions which occur in abundances from dominating sets [41]. To focus on the most common sequencing variants and to reduce runtimes, we first filtered to dominating set pieces that were present in at least 100 (16.5%) metagenomes; corncob fits a model to each dominating set piece, so it does not require abundance information for all pieces. We performed differential abundance testing using the `bbdml()` function using a likelihood ratio test with `formula = ~study_accession + diagnosis` and `formula_null = ~study_accession`. We estimated the number of k-mers in the quality controlled metagenome reads using ntcard and used this as the denominator. We performed Bonferroni p value correction and used a significance cut off of 0.05.

To analyze the results of dominating set differential abundance analysis, we combined the significantly differentially abundant piece information with the results from the mulitfasta query annotations, and with the ortholog annotations for the multifasta query genomes. We focused our analysis on KEGG orthologs. When a single gene was annotated by eggnog with multiple KEGG orthologs, we selected the first match. We identified the set of KEGG orthologs that were annotated among pieces that increased in abundance and pieces decreased in abundance in CD compared to nonIBD. To identify single copy marker genes within this set, we used the marker gene sequences with an average copy of one in [46]. We then selected at least one marker gene for each species, focusing on `rpl` sequences that were only annotated on two differentially abundant pieces in the graph (*Enterocloster clostridioformis, rplT; Ruminococcus_Bgnavus, rplQ Enterocloster clostridioformis_A, rpIO, rpIC; Enteroclostersp005845215, rpIO; Enterocloster bolteae, rplC*). We queried with the marker gene sequence that was used to perform the multifasta annotation and extracted the reads associated with that graph piece using `spacegraphcats extract_reads`. We then mapped the extracted reads back to the query sequence using `bwa mem` [69]. We visualized alignments in the Integrative Genomics Viewer [70] and selected reads that overlapped the same coordinates in the reference gene but that had different complements of single nucleotide polymorphisms. We BLASTed these reads using blastn against the NCBI nr database and found the best strain-level matches.

We next identified KEGG orthologs that were only annotated in the either the pieces that were increased or decreased in abundance in CD compared to nonIBD. We performed KEGG enrichment analysis using clusterProfiler `enricher` [71], using TERM2GENE as all KEGG orthologs with pathway mappings and with argument `maxGSSize = 500`. We considered pathways enriched to be enriched which had adjusted p values < 0 .05. Lastly, we searched for the presence of KEGG orthologs that quench reactive oxygen species using orthologs defined in [47].

### Searching for isolates that contained differentially abundant genomic sequences

To identify isolate genomes that contained sequences that were in CD, we searched the GTDB rs202 database. We generated FracMinHash signatures (k = 31, scaled = 2000) of differentially abundant sequences using sourmash `sketch`. We searched GTDB rs202 using sourmash `search`, using parameter `-max-containment`. We filtered results to only include isolate genome sequences (e.g., removed metagenome-assembled genomes) and selected the top match as the best match.

### Searching for metagenomes that contained differentially abundant genome sequences

We intersected FracMinHash signatures (*k* = 31, scaled = 2000) of differentially abundant sequences and query neighborhoods for each genome query, producing hashes that were differentially abundant and observed within each metagenome. We combined these hashes across diagnosis conditions (CD, UC, and nonIBD) and used the complexUpset R package to visualize the intersection size across conditions.

## Supplementary information

**Figure S1:**
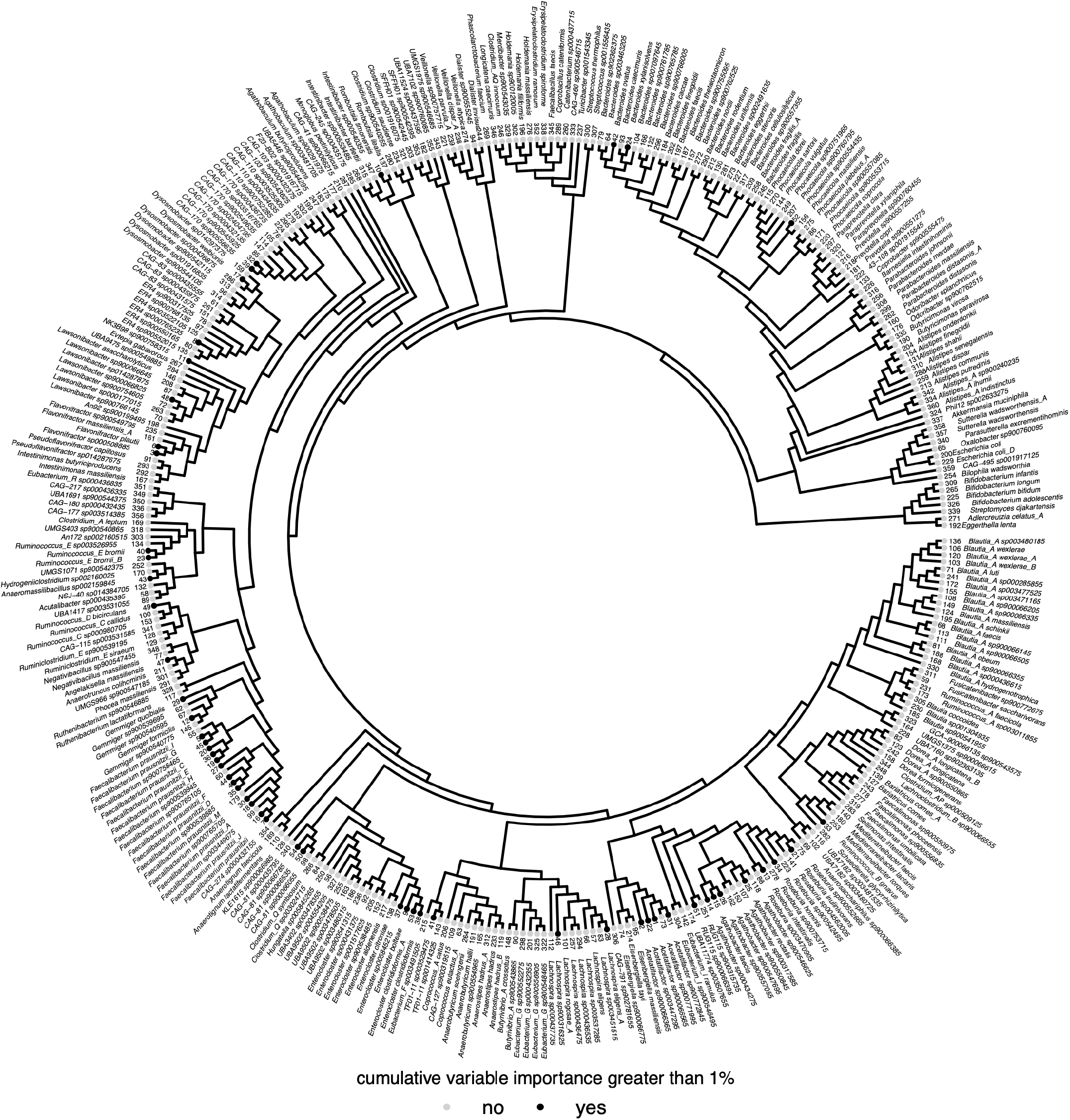
Phylogenetic tree of 360 bacterial species that were predictive of IBD subtype in all models. Tree was built from the GTDB rs202 tree with all tips except those represented by the 360 genomes removed. Tree tips are labelled by genomes that anchored at least 1 % of the normalized variable importance. The inner ring annotates the rank of the genomes, with the genome holding the most normalized variable importance across models ranked as 1. The outer ring is the species name within the GTDB database.

**Figure S2:**
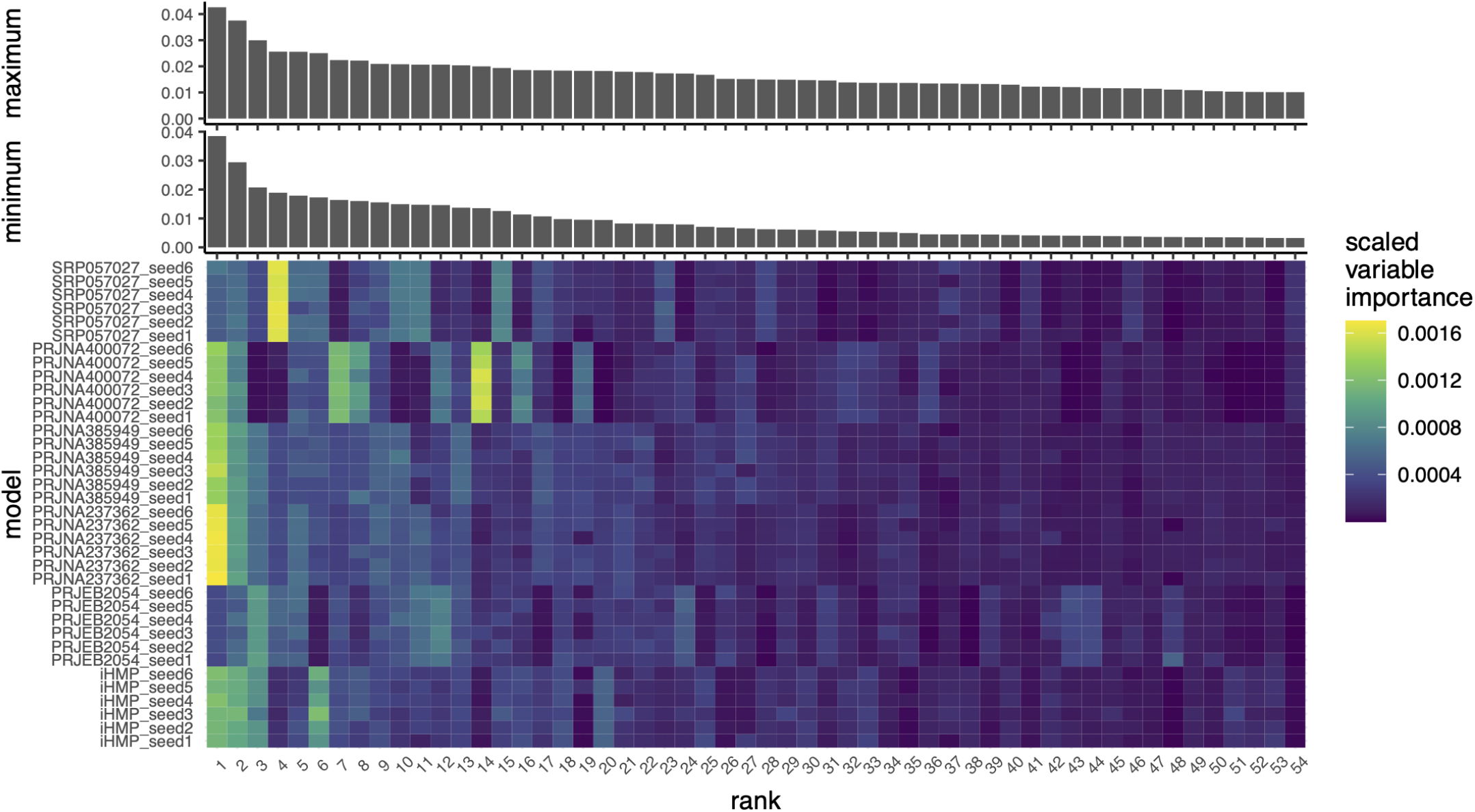
Fifty-four genomes are important across models and anchor the majority of variable importance. The bottom panel depicts a heat map of the scale variable importance contributed by k-mers that anchored to each of the top 54 genomes that were important for predicting IBD subtype. Models are labelled by the validation study and by the random seed used to build the model.Rank corresponds to the genome that anchored the most variable importance. Rank:species can be decoded using the tree in **Figure S1**. The top panels depict bar charts that correspond to the minimum (lower) or maximum (upper) variable importance a genome could anchor. The minimum variable importance was estimated following the sourmash gather algorithm, where each important k-mer was assigned to only one genome, and the genome it was assigned to was determined by a greedy winner-takes-all approach. Therefore, in the minimum bar chart, variable importance attributable to a k-mer was only summed once per k-mer, even if that k-mer occurred in multiple genomes. The maximum variable importance was estimated by allowing k-mers to be anchored to multiple genomes, so all k-mers were assigned to all possible genomes even if that meant a k-mer was assigned multiple times.

**Figure S3:**
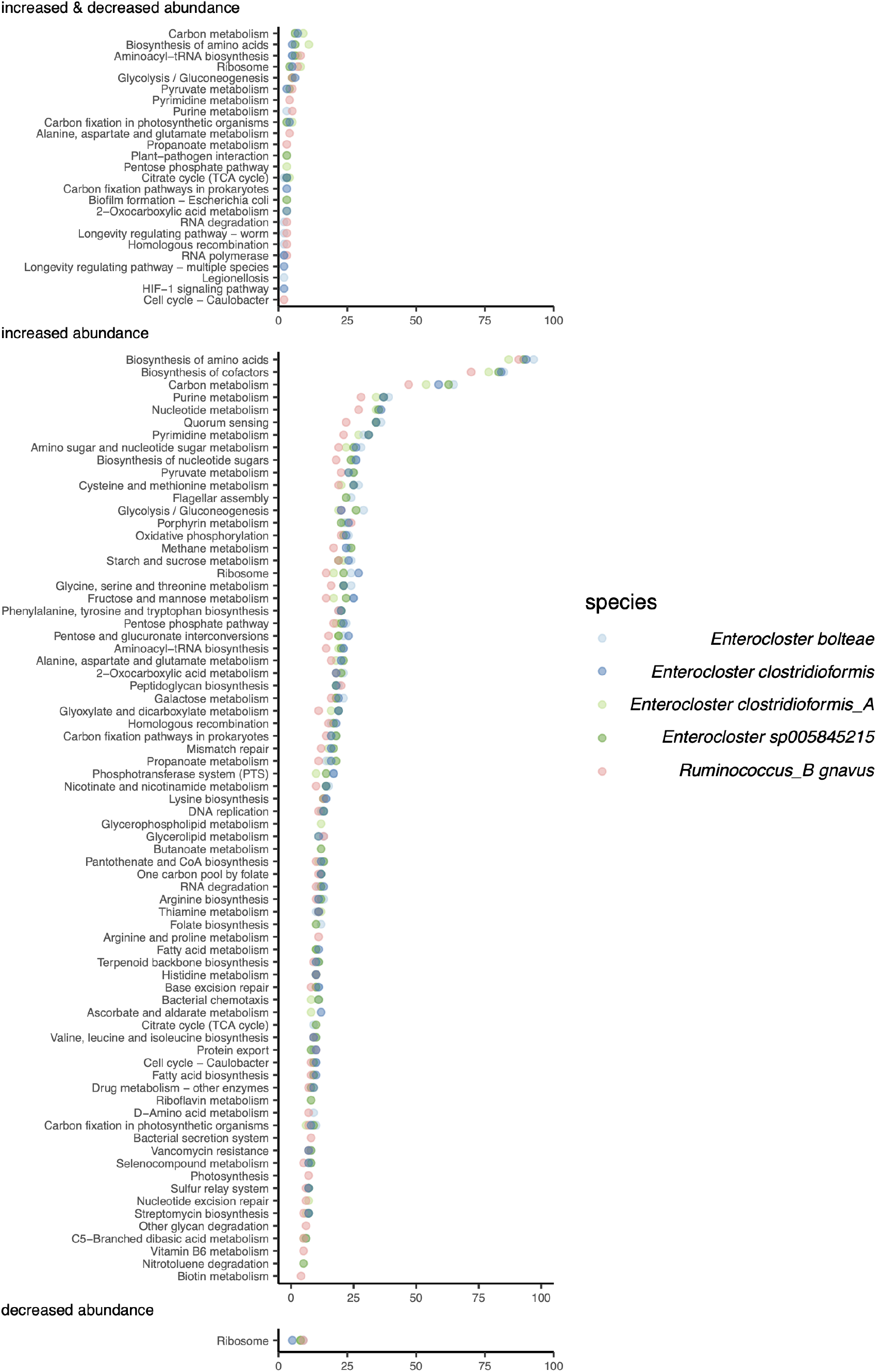
Pathways that were enriched among sets of differentially abundant sequences in CD compared to nonIBD. The x axis represents the number of orthologs identified in the pathway, while the y axis annotates the pathway. Top: Some dominating set pieces that significantly increased in abundance were annotated as the same KEGG orthologs as dominating set pieces that were significantly decreased in abundance. Many of these pathways encode core functions. Middle: KEGG pathway enrichment from KEGG ortholog annotations that were only observed in dominating set pieces that were significantly increased in abundance in CD. Bottom: KEGG pathway enrichment from KEGG ortholog annotations that were only observed in dominating set pieces that were significantly decreased in abundance in CD.

**Figure S4:**
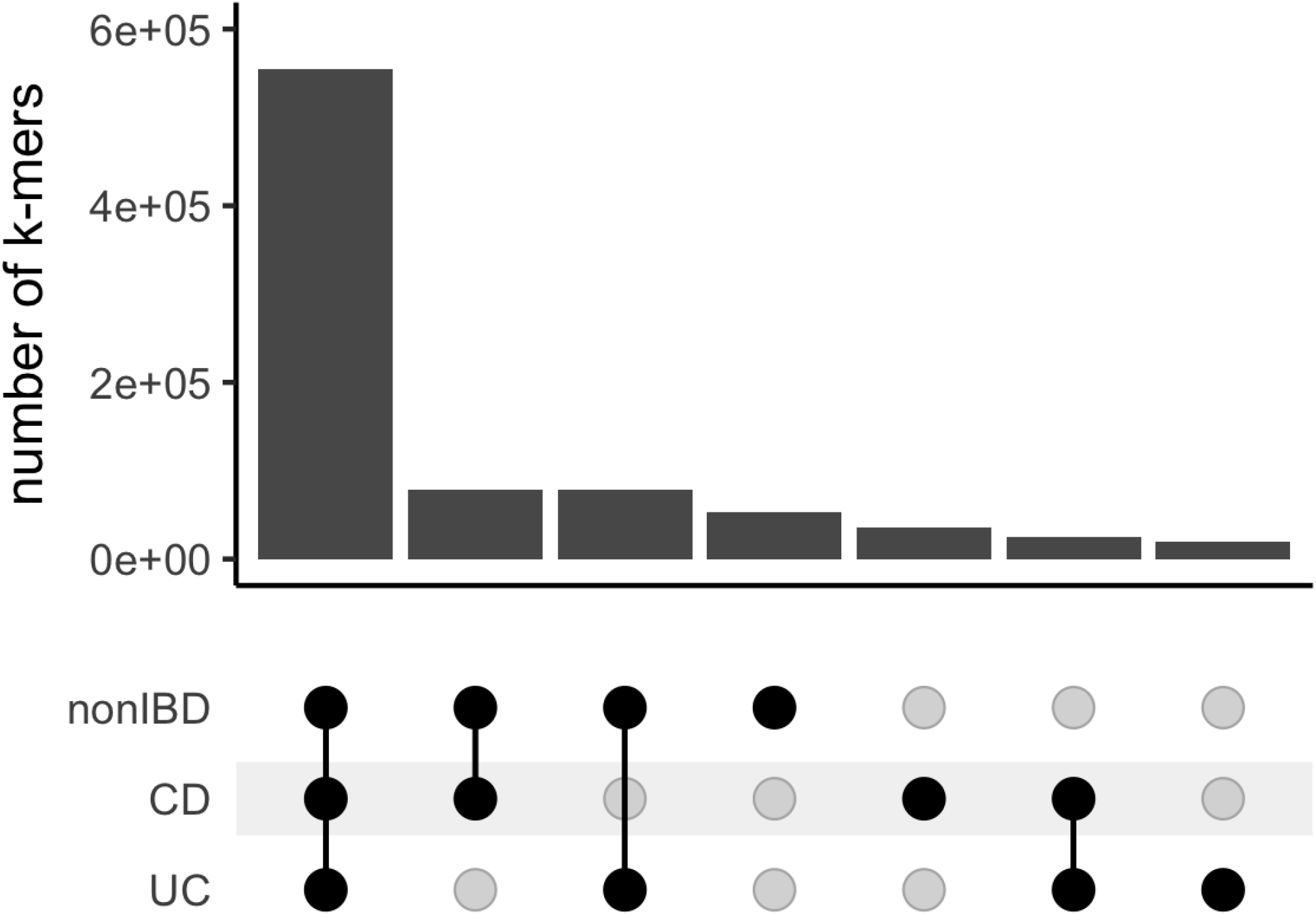
Most differentially abundant sequences occur in metagenomes of individuals diagnosed with CD, UC and non-IBD. Upset plot of k-mers that were decreased in abundance in CD and their occurrence in CD, UC, and nonIBD metagenomes. The bottom half of the plot highlights which diagnoses are included in each set, while the bar chart in the top half of the plot shows the number of k-mers that were observed in that set. The bar chart is colored by the metapangenome species graph in which the sequence was differentially abundant.

